# Island syndrome in the critically endangered Lord Howe Island cockroach *Panesthia lata*

**DOI:** 10.1101/2025.08.12.669983

**Authors:** Susannah K. Coady, Maxim W.D. Adams, Harley A. Rose, James A. Walker, Ian Hutton, Ros Gloag, Nathan Lo

**Affiliations:** School of Life and Environmental Sciences, University of Sydney, Sydney NSW 2006, Australia; Australian Government Department of Agriculture, Water and Environment, Canberra ACT 2601, Australia; Lord Howe Island Museum, Lord Howe Island NSW 2898, Australia

**Keywords:** island biology, island syndrome, animal behaviour, conservation, insects

## Abstract

Following island colonisation, organisms experience a unique array of selective pressures, giving rise to a somewhat predictable suite of morphological, demographic and ecological adaptations known as the “island syndrome”. Studies of the island syndrome have provided valuable insights into processes of speciation, community assembly, adaptive radiation and ecological release, alongside many others. However, to date, behavioural aspects of island adaptation have comparatively received little scientific attention, especially among invertebrates. In this study we examined the agonistic, courtship and aggregation behaviour of the endangered Lord Howe Island cockroach *Panesthia lata*, and compared these to its Australian sister species *Panesthia cribrata*. Behavioural assays revealed that while courtship behaviour was relatively stable across the two species, there was a significantly lower incidence of male agonism in *P. lata*. In concordence, analyses of nuclear single-nucleotide polymorphisms showed that *P. lata* forms large aggregations of unrelated individuals, unlike most *Panesthia* species, which maintain stable family groups. These results align with previous findings of relaxed intraspecific aggression in island mammals and reptiles, providing novel evidence of behavioural island syndrome in an invertebrate. We also found that courtship behaviour did not vary when *P. lata* interacted with conspecifics from the same or different populations, suggesting that individuals from different populations will readily interbreed. This is a promising outcome for the conservation of this critically endangered species, which currently spans a fragmentary range consisting of small, insular populations.

## 1. Introduction

Islands are valuable natural laboratories in which to study the process of evolution. Characteristics such as insularity, environmental unpredictability and low species diversity foster unique selective pressures (Covas, 2012; Losos & Ricklefs, 2009). As a result, many island taxa differ morphologically, ecologically, physiologically and demographically from mainland relatives, a phenomenon collectively termed the “island syndrome” (Baeckens & Van Damme, 2020). While the generality of island syndrome remains highly contested (reviewed by Jezierski et al., 2024), a number of adaptations appear to arise convergently across a broad swathe of taxa. For example, selection against flight has led to wing reduction in many insular birds and invertebrates (Leihy & Chown, 2020; Wright et al., 2016). Other widely reported adaptations include reduced reproductive output, higher life expectancy, heightened sexual dimorphism and potentially changes in body size (Adler & Levins, 1994; Baeckens & Van Damme, 2020; Benítez-López et al., 2021; Sen et al., 2014; Whittaker et al., 2023) but see (Biddick & Burns, 2021; Meiri et al., 2004, 2006).

Despite the adaptive significance of behaviour, there have been comparatively few examinations of the behavioural components of island syndrome. Island species often experience increased population density and ecological release, the combination of which appear to reduce boldness, aggression and intra-specific agonism (Baeckens & Van Damme, 2020). Similarly, the trend towards reduced fecundity may co-evolve with shifts in mating and parental care strategies (Covas, 2012). However, few studies have experimentally compared island species with mainland species, in particular for more complex traits such as behavioural syndromes (reviewed by Gavriilidi et al., 2022). Moreover, despite the fact that invertebrates are relatively over-represented in island ecosystems, research on island behaviour has overwhelmingly focused on vertebrates (reviewed by Gavriilidi et al., 2022; Meiri, 2007; New, 2008; Whittaker et al., 2023).

One potentially informative system in which to investigate the behaviour of island invertebrates is the Lord Howe Island Group (LHIG) of Australia. Located *ca*. 780 km northeast of Sydney, the LHIG comprises 28 islets and rocks, including Lord Howe Island (LHI) itself, with a total area of only 15 km^2^ (Green, 1993; Figure 1a). The islands support a diverse and highly endemic biota, which is overwhelmingly dominated by invertebrates (DECC, 2007). Surveys have reported over 1600 species of terrestrial invertebrate alone, with many more yet to be described (Cassis et al., 2003). While biogeographic studies are scarce, species appear to originate mainly from Australia, with contributions from New Zealand, New Caledonia, and other Pacific islands (Green, 1993). Small and remote islands such as the LHIG are thought to select most strongly for island syndrome (Whittaker & Fernández-Palacios, 2007), thus a comparison of an LHIG-endemic invertebrate with Australian relatives has great potential to reveal behavioural adaptations to island life.

**Figure 1.**
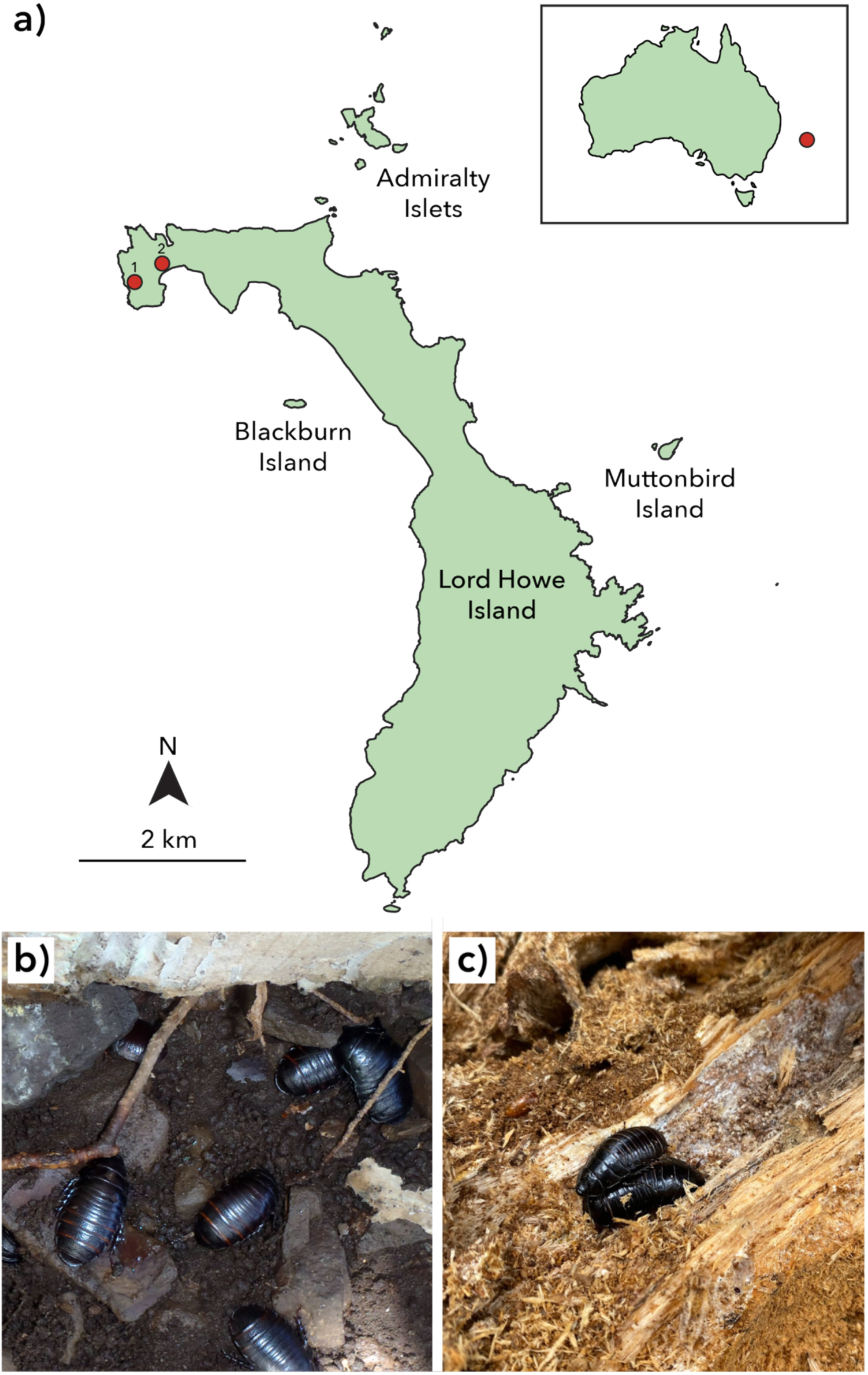
**a)** Map of the Lord Howe Island Group. Red dots denote the locations of the (1) North Head and (2) North Bay populations of *Panesthia lata*. Inset: position of the islands relative to mainland Australia. **b)** Large aggregation of *Panesthia lata* in soil on Blackburn Island. Photo: N. Carlile. **c)** Aggregation of *Panesthia cribrata* within a *Melalueca* sp. log in Bournda, New South Wales. Photo: M.W.D. Adams.

The Lord Howe Island cockroach *Panesthia lata* is a large, flightless insect endemic to the LHIG, estimated to have reached the islands *ca*. 2 Ma (Adams et al., 2025a). The species shows multiple ecological shifts relative to its congeners on the Australian mainland. The majority of *Panesthia* species reside within and feed upon decaying logs, aggregating in family groups that consist of one or both parents and several offspring. In the sister species of *P. lata*, *Panesthia cribrata sensu stricto* (*P. cribrata* South; Adams et al., 2025a) females likely remain within a single log for the duration of their lives, whereas males disperse at maturity (Rugg & Rose, 1984). In contrast, *P. lata* resides in shallow soil burrows beneath stones or logs, forming larger aggregations comprising multiple generations of adults and/or nymphs (up to *ca.* 15 individuals; M.W.D. Adams, pers. obs.). Individuals are also thought to leave the resident stone at night to forage for woody material and leaves (Carlile et al., 2018). While male *P. cribrata* readily engage in agonistic behaviours (O’Neill et al., 1987), it is unknown whether the higher aggregation densities of *P. lata* entail a reduction of intraspecific aggression, as is found in many island vertebrates. In addition, the kinship within and between its nesting aggregations has never been investigated with genetic data.

Behavioural study of *P. lata* will not only illuminate behavioural adaptations to island life, but will also augment a knowledge base dedicated to the species’ conservation. *Panesthia lata* is present on multiple islands of the LHIG, with the populations having been isolated by sea-level inundation for at least 10,000 years (Adams et al., 2025b). It was thought that population on LHI was locally extirpated by invasive black rats (*Rattus rattus*), leading the species to be listed as Endangered by the NSW Threatened Species Scientific Committee (2004), followed by a recent reassessment as Critically Endangered (NSW Threatened Species Scientific Committee, 2025). However, following the successful eradication of rodents in 2019, a relict population was discovered in LHI’s North Bay. The authors have shown that this population, and those on the smaller islands, are highly inbred, which may elevate the species’ extinction risk (Adams et al., 2025b). An understanding the ecology and behaviour of the different populations will contribute to any future conservation actions.

In this study, we undertake further surveys of LHI and examine the behaviour of *P. lata*. By examining social, agonistic and courtship behaviours in multiple populations across the LHIG, and comparing these to *P. cribrata*, we aim to: 1) determine whether *P. lata* displays behavioural changes consistent with the island syndrome; and 2) test whether courtship and agonistic behaviours vary between populations of *P. lata*. We also analyse a panel of nuclear single-nucleotide polymorphisms (SNPs) to 3) characterise patterns of genetic relatedness within nesting aggregations of *P. lata*. Our results reveal complex behavioural shifts following island colonisation, and provide further information to guide the management of this critically endangered species.

## 2. Material and Methods

### 2.1. Habitat surveys and specimen collection

Over four separate expeditions between 2022–2025, we opportunistically examined suitable habitats across LHI for the presence of *P. lata*. Informed by the species’ habitat preferences on Blackburn Island (Carlile et al., 2018; Rose, 2003), we focused our search on sites with moist soil, dense canopy and abundant ground cover consisting of decaying logs and/or stones, with a particular emphasis on areas in the vicinity of Banyan trees (*Ficus macrophylla* var. columnaris). In total, we surveyed five different regions of the island: Old Settlement, Stevens Reserve, Valley of Shadows, Soldiers Creek, and along parts of Lagoon Road.

At a given location, we searched for cockroaches by overturning leaf litter and any large (> *ca.* 20 cm) stones, as well as breaking any large (> *ca.* 10 cm diameter) logs. All surveys were undertaken during the day. No cockroaches or traces were found at all but two sites. In addition to a previously discovered relict population in North Bay (Adams et al., 2025b; Figure 1), individuals of *P. lata* were detected by Toby Kovacs, SC, and NL for the first time at a site on the island’s North Head (Figure 1) on May 24th, 2023. At this location, both adult and juvenile cockroaches were found sheltering beneath stones directly under the canopy of a single *F. macrophylla* tree. Despite its geographic proximity to North Bay (*ca.* 400 m), we consider this a separate population due to the lack of suitable intervening habitat. Further ecological observations, based on a transect conducted on February 20^th^ 2025, are outlined in the Supplementary Material.

In May 2023, individuals of *P. lata* were collected from three separate populations for our behavioural assays: Blackburn Island (31.534°S 159.060°E), North Bay, LHI (31.517°S 159.042°E) and North Head, LHI (31.517°S 159.038°E). Prior to collection, we surveyed both relict populations to ensure a sufficient population density. Cockroaches were collected in “aggregations”, which we defined as all individuals under a single rock or log. We collected six aggregations from Blackburn Island, nine from North Bay and six from North Head We collected specimens of *P. cribrata* from two separate localities in New South Wales: Harold Reid Reserve, Sydney (33.796°S 151.212°E) and Monga State Conservation Area, Braidwood (35.445°S 149.900°E). Collections were undertaken between April and August 2023. We collected discrete aggregations as follows: decaying logs, branches or tree trunks were overturned, and any visible cockroaches were collected; logs were then prised open to search for resident galleries containing additional cockroaches. Individuals in connected galleries within the same log, or in soil < 2 m from the galleries, were considered a single aggregation; while individuals in unconnected galleries or separated by > 2 m were considered a separate aggregation. In total, five aggregations of were collected from Sydney and nine were collected from Braidwood. Cockroaches from both species were cultured at ambient temperature in their aggregation groups, and provided with coconut coir as substrate. Water and wood chips were provided *ab limitum*. To permit identification in behavioural assays, each adult was marked with a uniquely coloured and numbered acrylic marker indicating its population and aggregation, which was glued onto the metanotum.

Finally, to investigate pairwise kinship within and between nesting aggregations of *P. lata*, we collected samples for SNP data generation in July 2022. We collected individuals from under four separate rocks on Blackburn Island, chosen such that each rock was more than 5 m from all others in the sample (comfortably exceeding the expected foraging range of *P. lata* in a single night, ≤ 1.12 m; Carlile et al., 2018). Care was taken to collect every individual present under each rock, and samples were stored in 96–100% ethanol prior to DNA extraction. The number of individuals in each aggregation ranged between *n* = 3–9. In addition, we retrieved putative unrelated individuals (i.e. all from separate aggregations) from Blackburn Island, Roach Island and North Bay, LHI to include as baseline reference points in our analyses.

Details of their collection can be found in Adams et al. (2025b).

### 2.2 Aggregation size, morphometrics and nesting behaviour

We recorded the total number of individuals in each collected aggregation and measured the body length of all available adults of *P. lata* and *P. cribrata*. Live measurements were taken with a ruler to the nearest millimetre, measured from the anteromedial margin of the pronotum to the posteromedial margin of the supraanal plate.

We tested whether aggregation size varied across all five populations (from both species) using a Kruskal-Wallis test in R. We also used a Kruskal-Wallis test to determine whether body length differed between populations, performed separately for males and females. Where significant overall differences were found, we performed post-hoc individual comparisons using Dunn tests (package *dunn.test* v.1.3.6; Dinno, 2017) with Bonferroni corrections.

To further investigate nesting behaviour and movement by *P. lata*, we performed two marking experiments on Blackburn Island, on May 19^th^ 2023 and September 14^th^ 2024. In each case, all individuals under each of 6 rocks, spaced >5 m from each other under the single *Ficus* tree on the island were marked with a different colour of nail polish across their metanotum and dorsal abdomens (an area of ∼3 cm^2^ in adults). The rocks under which they were found were marked, and individuals were returned to their respective rocks. On the evenings of May 23^rd^ and September 17^th^, the six marked rocks, as well as rocks within 1-2 metres of the marked rocks, were checked for the presence of *P. lata*, with each individual carefully checked for the presence of nail polish.

We also conducted observations of *P. lata* under Blackburn Island’s *Ficus* tree at 9 pm on May 23^rd^ 2023. This was to confirm that the species is nocturnal, and whether it forages in the open, or remains under leaf litter. Initial general observations were made prior to conducting a survey where three field workers walked from one side of the *Ficus* to the other (*ca.* 20 m), recording the life stage (adult or nymph) and sex of any *P. lata* seen on the ground (*ca.* 20 minutes of observation).

### 2.3. Behavioural assays

We examined behaviour using a custom-built arena. This consisted of 3 *ca.* 5×5 cm chambers, connected by corridors *ca.* 2 cm wide (Figure 2), carved into a pine-wood block to a uniform depth of *ca.* 1.5 cm. The arena was covered by a sheet of translucent Perspex to maintain humidity. Chambers were furnished with peat moss, as well as small pieces of wood or *Ficus* leaves for *P. cribrata* and *P. lata*, respectively.

**Figure 2.**
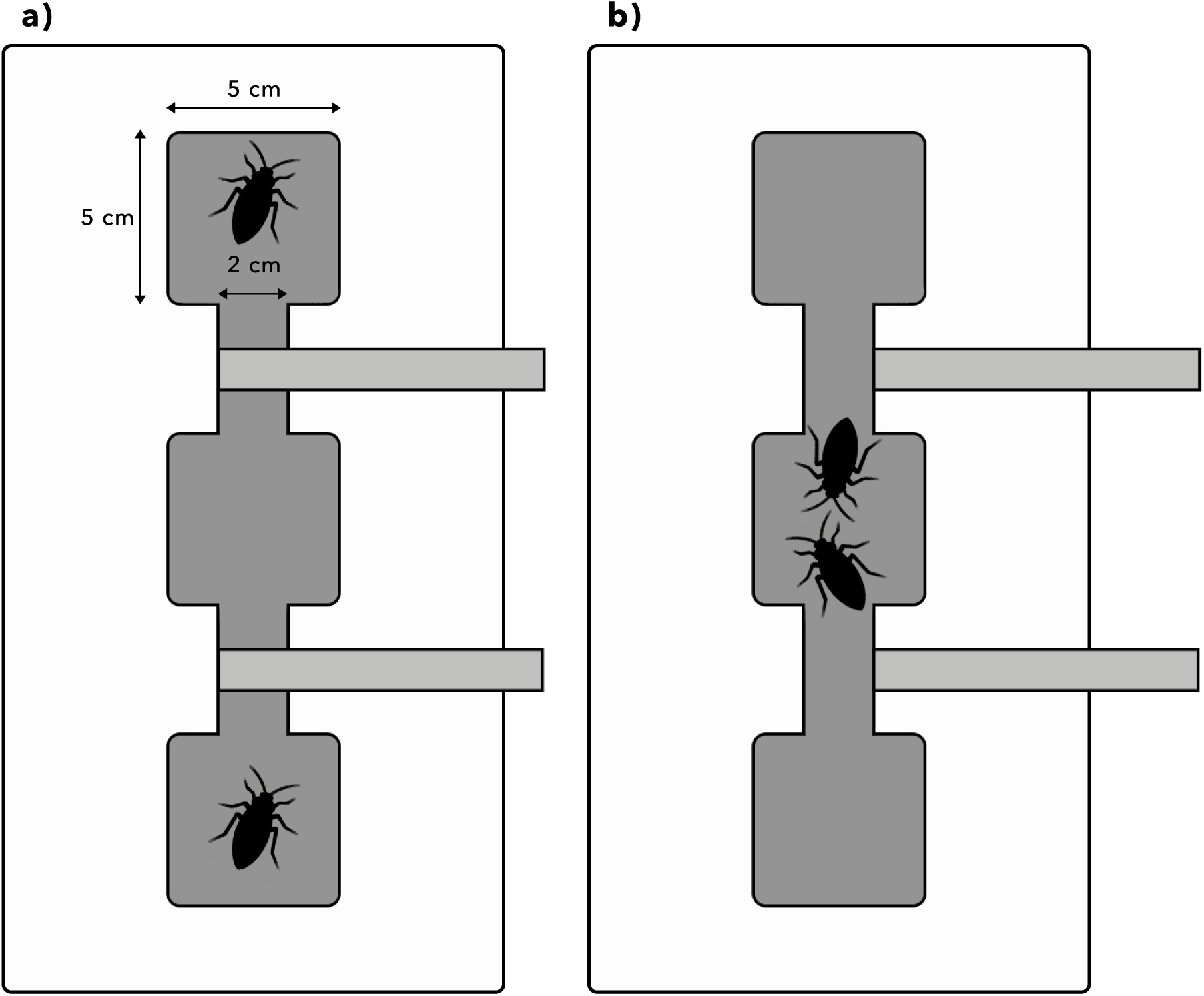
Top-down schematic diagram of a behavioural arena. Dark shaded region represents three chambers with intervening corridors. Light shaded rectangles represent wooden rods, which are slotted into carved grooves and can be moved left or right (in top-down view). **a)** Prior to the behavioural assay, wooden barriers prevent interaction. **b)** Barriers have been withdrawn to initiate the assay and permit interaction between cockroaches.

We investigated agonism and courtship by conducting behavioural assays with pairs of adult cockroaches. In each assay, two marked cockroaches were placed in opposite chambers of the arena, initially with retractable wooden barriers blocking the corridors to prevent their exploration (Figure 2a). These were left undisturbed in a darkened cabinet for 12 hours to acclimate. To begin the assay, the barriers were manually withdrawn and the cockroaches’ interactions were recorded for *ca.* 8 hours using an iPhone camera. As *Panesthia* species are nocturnal and photophobic (O’Neill et al., 1987), each assay was initiated at approximately 22:00, and the arena was illuminated using red LED lighting. Agonistic assays consisted of male-male or female-female pairings, while courtship assays were conducted using male-female pairs (Table 1).

**Table 1:**
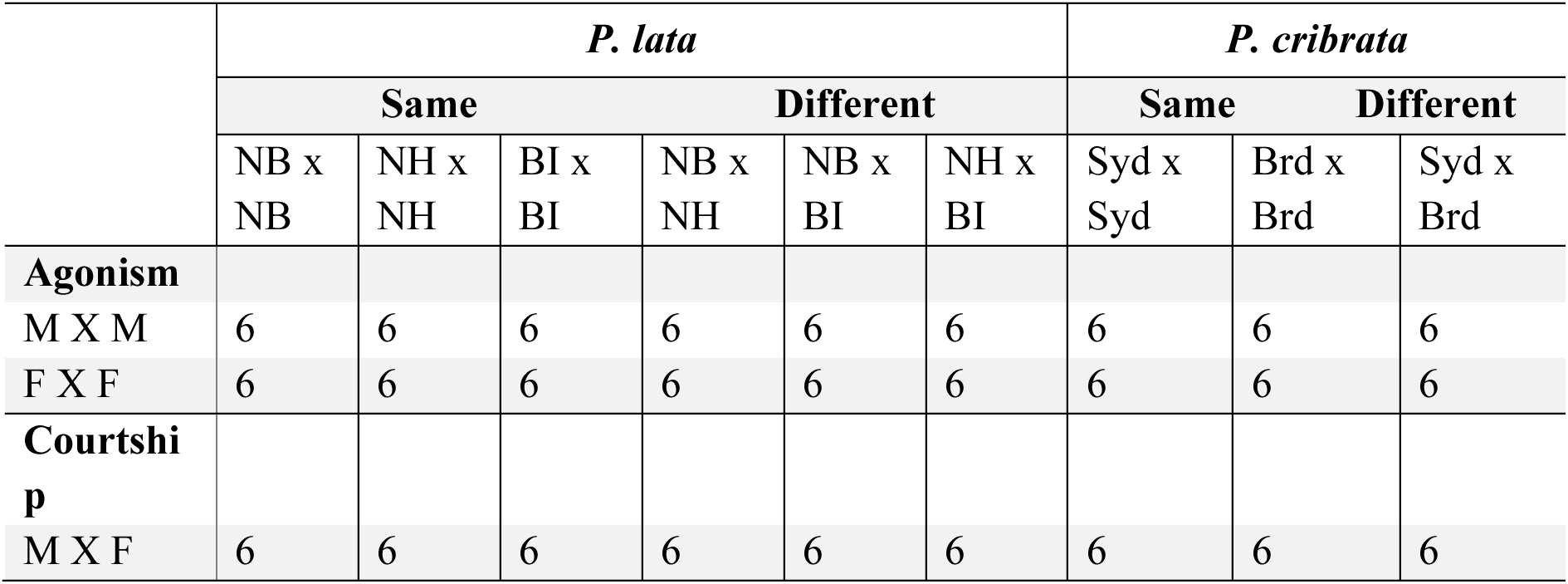
Nine treatment categories used to assess agonism and courtship behaviours in *P. lata* (three populations: BI = Blackburn Island, NB = North Bay, NH = North Head) and *P. cribrata* (two populations: Syd = Sydney, Brd = Braidwood). Six replicates were performed for each category. Treatments assessed the behaviours of *Panethsia* when interacting with members of their own populations, or conspecifics from different populations, and when interacting with either members of the same or different sexes.

Agonistic and courtship assays were conducted separately for *P. lata* and *P. cribrata*. For each species, we used a fully crossed design, whereby we tested behaviour using pairs of cockroaches from within each population and each combination thereof (Table 1). For intra-population assays, the two cockroaches were taken from different aggregations. We conducted six replicates of each of the nine population pairings, for all three sex pairings (male-male, female-female and male-female).

Videos were viewed and scored using QuickTime Player v.10.5 (Apple Computer, USA). Primarily following O’Neill et al. (1987), we recognised 13 different behaviours across three behavioural categories: neutral, agonistic or courtship (Table 2). For each video, we recorded the number and types of behaviours, the number of discrete interactions, the duration of each interaction, and the behavioural category of each interaction (neutral, agonistic or courtship). We considered an interaction to have begun when both cockroaches had > 50% of their body length within the same or adjacent chamber or corridor. We considered the interaction to have ended once the cockroaches had > 50% of their body length in separate chambers for over 60 seconds. The behavioural category of each interaction was assigned based on the modal behaviour.

**Table 2:**
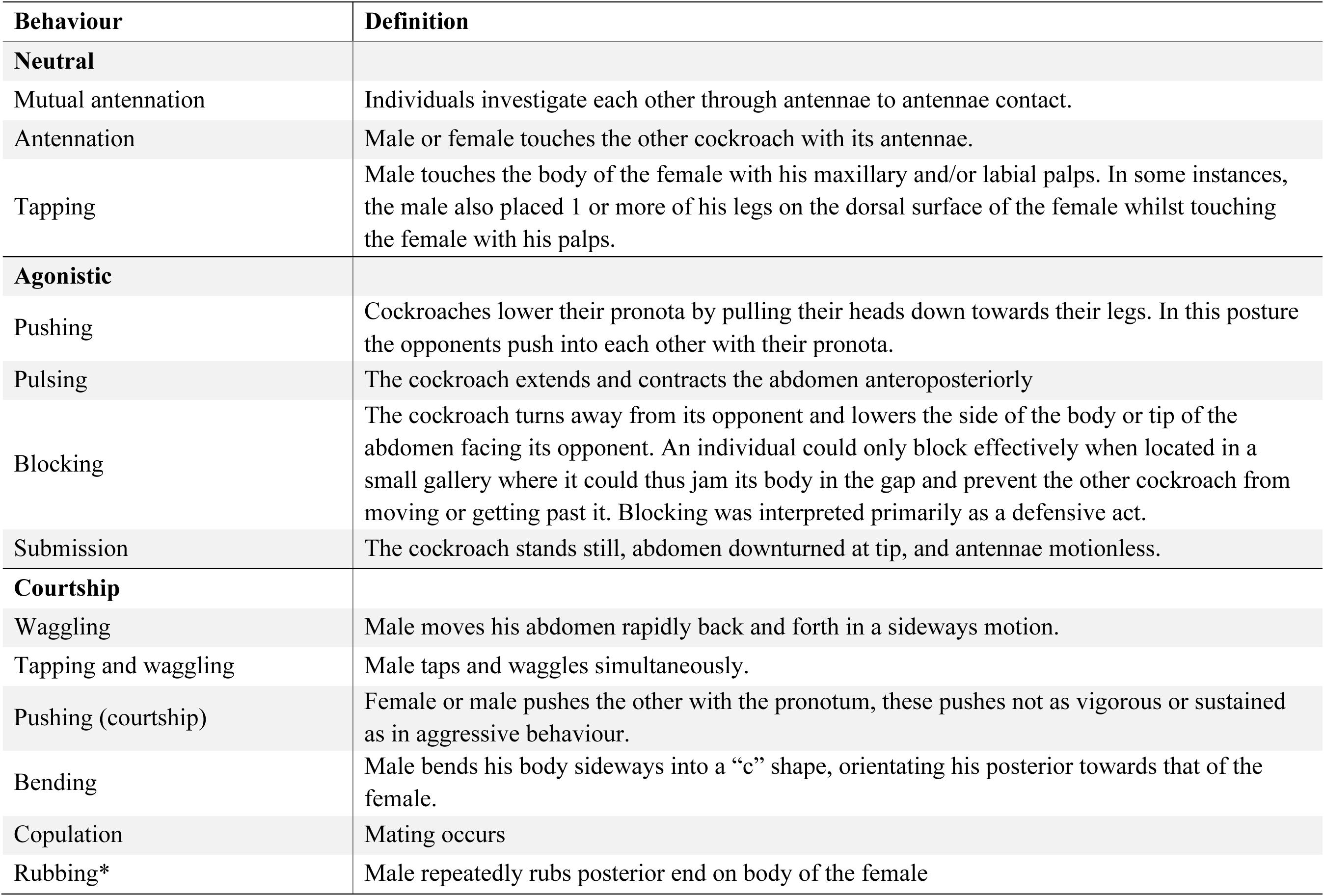
Categories of *Panesthia* behaviours (following O’Neill *et al*., 1987). Asterisk (*) indicates a newly defined behaviour.

We quantified behaviour using four outcomes: the number of agonistic/courtship interactions per hour, the proportion of agonistic/courtship interactions from the total number of interactions, the average duration of agonistic/courtship interactions, and the intensity of agonistic/courtship interactions. Intensity was scored as the number of different behaviours per interaction. After confirming data normality and homoscedasticity, we used generalised linear models in R to test whether these outcomes varied between species or between population pairings (whether the pairing comprised cockroaches of the same or different populations), and to check for interactions between these factors.

Second, for both agonistic and courtship interactions, we quantified the number of times each specific behaviour was performed per interaction (Table 1). We then tested whether the frequencies and types of behaviour varied between species and population pairings using a two-way multivariate analysis of variance (MANOVA) in R. Where significant differences or interactions were found, we undertook post-hoc tests examining variations in individual behaviours using two-way ANOVAs with Bonferroni corrections.

### 2.4. SNP data generation and analysis

We undertook genetic analyses using the four nesting aggregations of *P. lata* collected on Blackburn Island, as well as the putative unrelated individuals from Blackburn Island, Roach Island and North Bay (*n* = 67 individuals in total). No North Head samples were included in genetic analyses as the population was discovered subsequent to DNA sequencing. For the majority of samples, DNA extractions were carried out in-house, using a DNeasy Blood and Tissue Kit (Qiagen, Germany) per manufacturer’s instructions. We checked DNA purity and concentration using a NanoDrop 2000 spectrophotometer (Thermo Fisher Scientific, USA) and a Qubit 2.0 fluorometer (Invitrogen, USA), respectively. For a minority of samples, DNA extraction was instead outsourced to the Australian National Insect Collection, Canberra (see Adams et al., 2025a for details; Supplementary Table S2).

Reduced-representation sequencing and SNP data generation were carried out by Diversity Arrays Technology (DArT), Canberra. A subset of the data was previously analysed by Adams et al. (2025b), and sequencing methods are described in detail therein. In summary, DNA was digested with the *Pstl* and *Hpall* restriction enzymes, sequenced using a Novaseq 6000 S1, and processed through the proprietary DArT analytical pipeline.

Data were filtered chiefly using the R package *dartR* v.2.7.2 (Gruber et al., 2018). We first filtered for data quality and completeness, based on reproducibility (100%), minor allele frequency (MAF > 0.025; based on 3/2*N*) and call rate (> 0.8, calculated for both individuals and loci). This “quality-filtered” dataset comprised 11,446 SNPs from 62 individuals. We then produced a more stringently filtered dataset of putatively unlinked and neutral SNPs.

Monomorphic loci were removed, and we retained one randomly selected SNP per sequence tag (removed “secondaries”). We also removed loci out of Hardy-Weinberg equilibrium based on the exact test (alpha value = 0.01), calculated with the mid-p method. Finally, we confirmed the absence of any loci potentially under selection (based on elevated *F*_ST_ values) using BayeScan v.2.1 (Foll & Gaggiotti, 2008) under default settings. The “neutral” dataset comprised 8,006 SNPs from 62 individuals. Average coverage for both data sets was *ca.* 12 ×.

We used two approaches to explore the relatedness of individuals within and between the Blackburn Island aggregations. First, we conducted kinship analyses using the quality-filtered SNP panel, with individuals from Blackburn Island, Roach Island and North Bay included as putative unrelated reference points. We identified any closely related pairs (parents, grandparents or siblings) using the R package *Sequoia* v.2.11.2 (Huisman, 2017). In addition, we calculated individual pairwise kinship coefficients using maximum likelihood in the R package *SNPRelate* v.1.33.0 (Zheng, 2013). These values were averaged within each nesting aggregation (excluding self-kinship), as well as within the remaining, independent individuals from Blackburn Island.

Second, we visualised genetic structure within and between nesting aggregations. We pruned the dataset to include only the four Blackburn Island aggregations, and re-filtered the raw SNPs as above (producing a quality-filtered panel of 8,926 SNPs and a neutral panel of 6,471 SNPs, both from 24 individuals). Using the quality-filtered SNPs, we conducted a principal coordinates analysis (PCoA) of individual allele frequencies in *dartR*. Using the neutral SNPs, we analysed population structure using a sparse non-negative matrix factorization algorithm (sNMF) in the R package *LEA* v.2.0 (Frichot & François, 2015). We searched for the optimum number of genetic clusters between *K* = 1 to *K* = 8, with 10 replicates for each *K* value.

## 3. Results

### 3.1. *Nesting ecology of* Panesthia lata

In 2023, 33 wild individuals were marked with nail polish on Blackburn Island, across 6 separate aggregations. When the location was re-surveyed after four days, only a single marked individual was re-captured, found under a separate rock *ca.* 20 cm from the site of its first capture. Based on pilot studies, we cannot exclude the possibility that some markings were removed by abrasion.

However, all six rocks also sheltered a different number of adult males, females, and nymphs when reinspected after the four days, suggesting that there had been movement of almost all individuals to and from rocks. When repeated in September 2024, a total of 32 individuals across 6 rocks were originally marked. Upon re-inspection, no marked individuals were found under or near four of the six marked rocks. One rock, under which 1 male, 1 female, and 2 nymphs were found and marked, was found 3 days later to 1 marked individual (1 female), and a further 2 marked nymphs plus one unmarked individual, were found under a second nearby rock (∼30 cm from the original marked rock). A second of the original 6 rocks, under which 4 females, 1 adult, and 1 nymph were originally marked, was found to have one marked female 3 days later.

No cockroaches were detected outside rocks during the daytime at all sites examined; however, during a short observation period commencing 9 pm on May 23^rd^ 2023, adults were seen moving in the open (under cover of darkness). During the 20-minute night-time transect, 7 adult males, 10 adult females and no nymphs were sighted.

### 3.2. Aggregation size and morphometrics

The total number of individuals aggregating either under rocks (*P. lata*) or within rotten wood (*P. cribrata*) varied significantly between populations (χ²_4_ = 11.746, *p* = 0.019; Figure 3a). Specifically, aggregations were larger on Blackburn Island than in North Bay (post-hoc *p* = 0.019).

**Figure 3.**
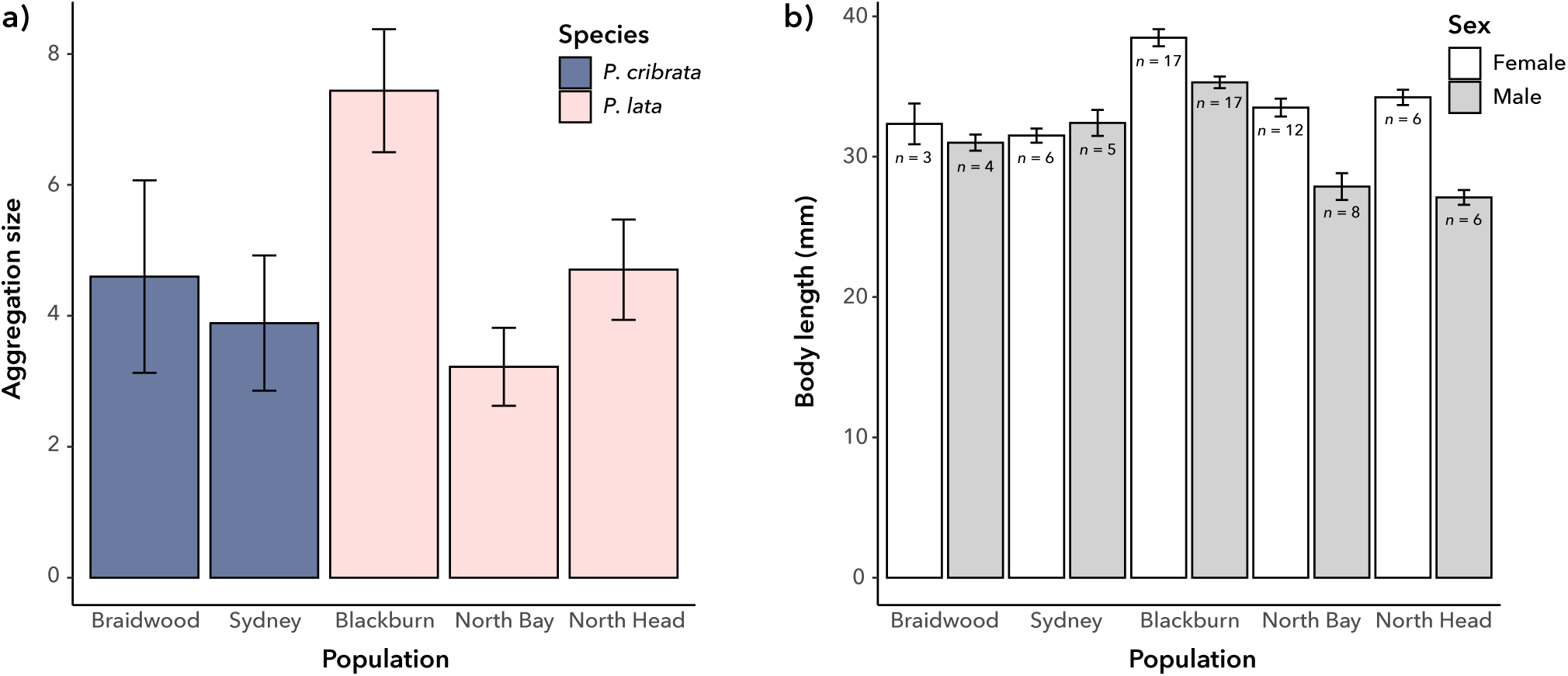
Comparisons of five populations of *P. cribrata* and *P. lata*. **a)** Mean aggregation size (+/- standard error). **b)** Mean body length (+/- standard error) of adult males and females. Bars are labelled with sample size (*n*).

Body length varied significantly between populations among both females (χ²_4_ = 30.139, *p* < 0.001; Figure 3B) and males (χ²_4_ = 34.842, *p* < 0.001; Figure 3b). Post-hoc comparisons indicated that females from Blackburn Island were longer than those from all other populations (*p* ≤ 0.015 for all), while males from Blackburn Island were longer than those from North Bay and North Head (*p* ≤ 0.001 for both).

### 3.2. Agonistic behaviour

We found no significant differences in all four measures of agonistic behaviour among female cockroaches, irrespective of species or population pairing, with no significant interactions. In contrast, male agonistic behaviour varied significantly between *P. cribrata* and *P. lata* (Figure 4). The number of agonistic interactions per hour (*t =* 3.542, *p* < 0.01), the proportion of interactions with agonism (*t* = 2.156, *p* = 0.0359), and the intensity of agonistic interactions (*t* = 2.537, *p* = 0.0143) were all significantly higher in *P. cribrata* (Figure 4a,b,d). However, the average duration of agonistic interactions was not significantly different between the species (*t* = 1.969, *p* = 0.0545; Figure 4c). None of the outcomes varied significantly based on population pairing, nor were there significant interactions between species identity and population pairing.

**Figure 4.**
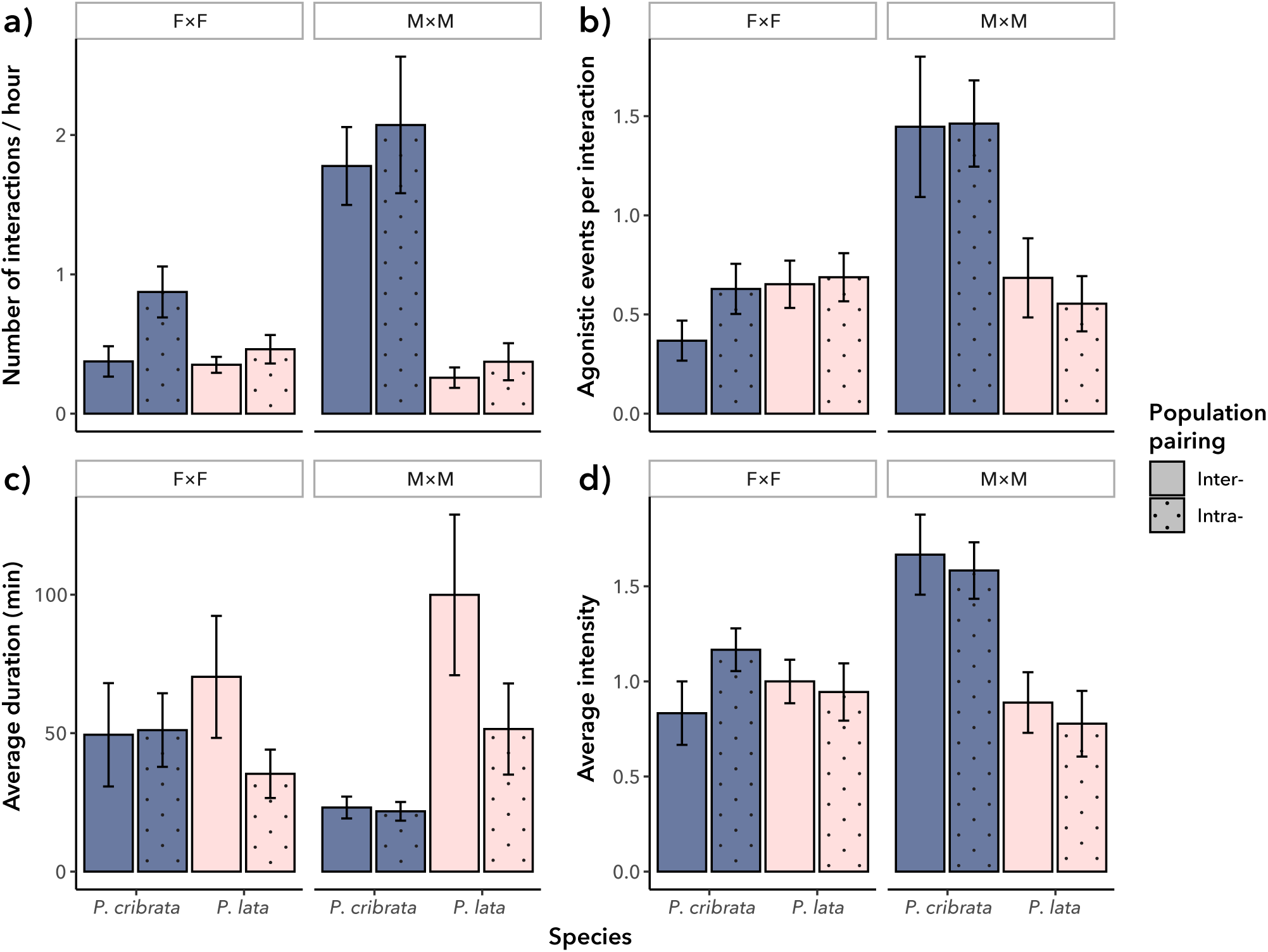
Agonistic behaviour in *P. cribrata* and *P. lata* (mean +/- standard error); F×F = female pariings, M×M = male pairings. **a)** Average number of agonistic interactions per hour. **b)** The proportion of the total number of interactions that were agonistic. **c)** The average duration of each agonistic interaction. **d)** The average intensity of each agonistic interaction.

The frequencies of different types of agonistic behaviour varied significantly between male *P. cribrata* and *P. lata* (MANOVA: F_1,47_ = 17.406, *p* < 0.01; Figure 5). Specifically, there was a significantly higher incidence of blocking in *P. cribrata* (F_1,50_ = 68.953, *p* < 0.01). Behavioural frequencies did not vary significantly between population pairings (F_1,47_ = 0.245, *p* = 0.911), nor was there a significant interaction between species and population pairing (F_1,47_ = 0.884, *p* = 0.481). For female agonistic behaviours, we found a significant interaction between species and population pairing (MANOVA: F_4,47_ = 8.475, *p* < 0.01; Figure 5). This was specifically driven by a significant interaction between species and population pairing for submission rate (F_1,50_ = 18.240, *p* < 0.01), whereby submission only occurred in intra-population pairings of *P. cribrata*.

**Figure 5.**
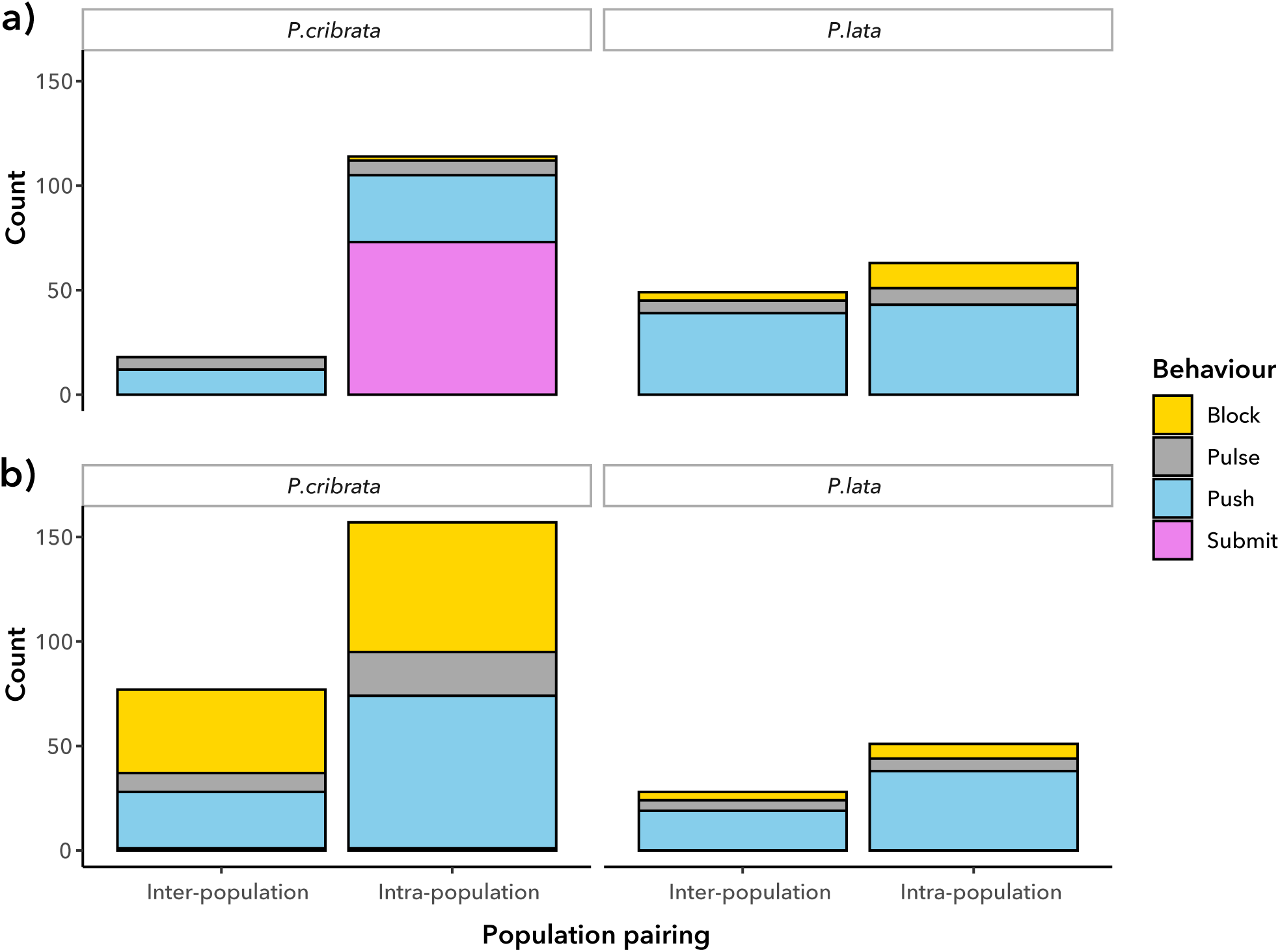
Proportions of individual behaviours in agonistic interactions. **a)** Behaviours in female-female interactions. **b)** Behaviours in male-male interactions.

**Figure 6.**
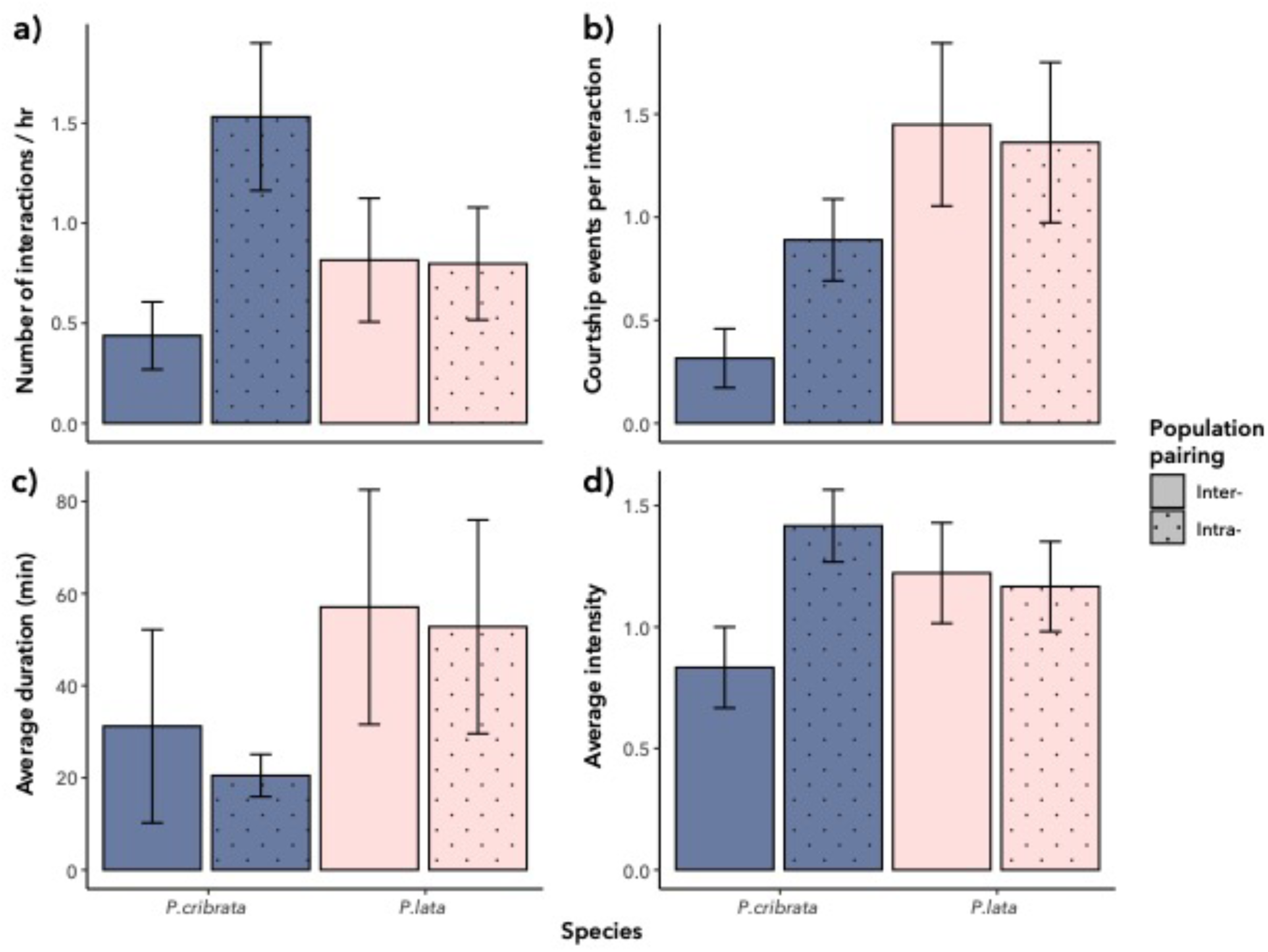
Courtship behaviour in *P. cribrata* and *P. lata* based on observation of male-female pairs (mean +/- standard error). **a)** Average number of courtship interactions per hour. **b)** The proportion of the total number of interactions that involved courtship. **c)** The average duration of each courtship interaction. **d)** The average intensity of each courtship interaction.

### 3.3. Courtship behaviour

None of the four measures of courtship behaviour varied significantly between species or population pairings, and we found no significant interactions between these factors. However, the frequencies of different courtship behaviours did vary significantly between *P. cribrata* and *P. lata* (F_1,45_ = 10.246, *p* < 0.01), but not based on population pairings (F_1,45_ = 1.032, *p* = 0.418).

There was no significant interaction between these factors (F_1,45_ = 2.081, *p* = 0.0743). *Panesthia cribrata* was found to perform the tap and waggle behaviour significantly more frequently than *P. lata* (F_1,50_ = 24.370, *p* < 0.01).

Finally, we observed courtship behaviours in six male-male trials of *P. lata*. The behaviours were all performed by males from Blackburn Island, when paired with individuals from Blackburn Island (*n* = 1 trial), North Bay (*n* = 3 trials) and North Head (*n* = 2 trials).

### 3.4. Kinship and genetic structure

Kinship estimates were conducted using the full dataset, consisting of both the Blackburn Island aggregations and putative unrelated individuals from Blackburn Island, North Bay and Roach Island. Pedigree analysis with *Sequoia* did not detect any parent, sibling or grandparent relationships among the Blackburn Island aggregations. In concordance, pairwise kinship analyses did not detect any close relatives (*k* > 0.125) within any of the aggregations. Average pairwise kinship within each aggregation was very similar to the average pairwise kinship between the remaining individuals from Blackburn Island (Table 3).

**Table 3.** Average pairwise kinship (excluding self-kinship) within and between aggregations of *P. lata* on Blackburn Island. Values calculated in *SNPRelate*. *N* = sample size.

| Group | $N$ | Average pairwise kinship |
| --- | --- | --- |
| Aggregation 1 | 3 | 0.0650 |
| Aggregation 2 | 3 | 0.0597 |
| Aggregation 3 | 9 | 0.0625 |
| Aggregation 4 | 9 | 0.0636 |
| Separate aggregations | 11 | 0.0636 |

Analyses of genetic structure were undertaken using only the four nesting aggregations from Blackburn Island. No structure was evident based on PCoA (Figure 8a). Similarly, clustering analysis with sNMF optimised at *K* = 1, indicating an absence of genetic structure (Figure 8b). Visualisations of structure and admixture at higher *K* values did not reveal any patterns corresponding to the four aggregations (Supplementary Figure S2).

**Figure 7.**
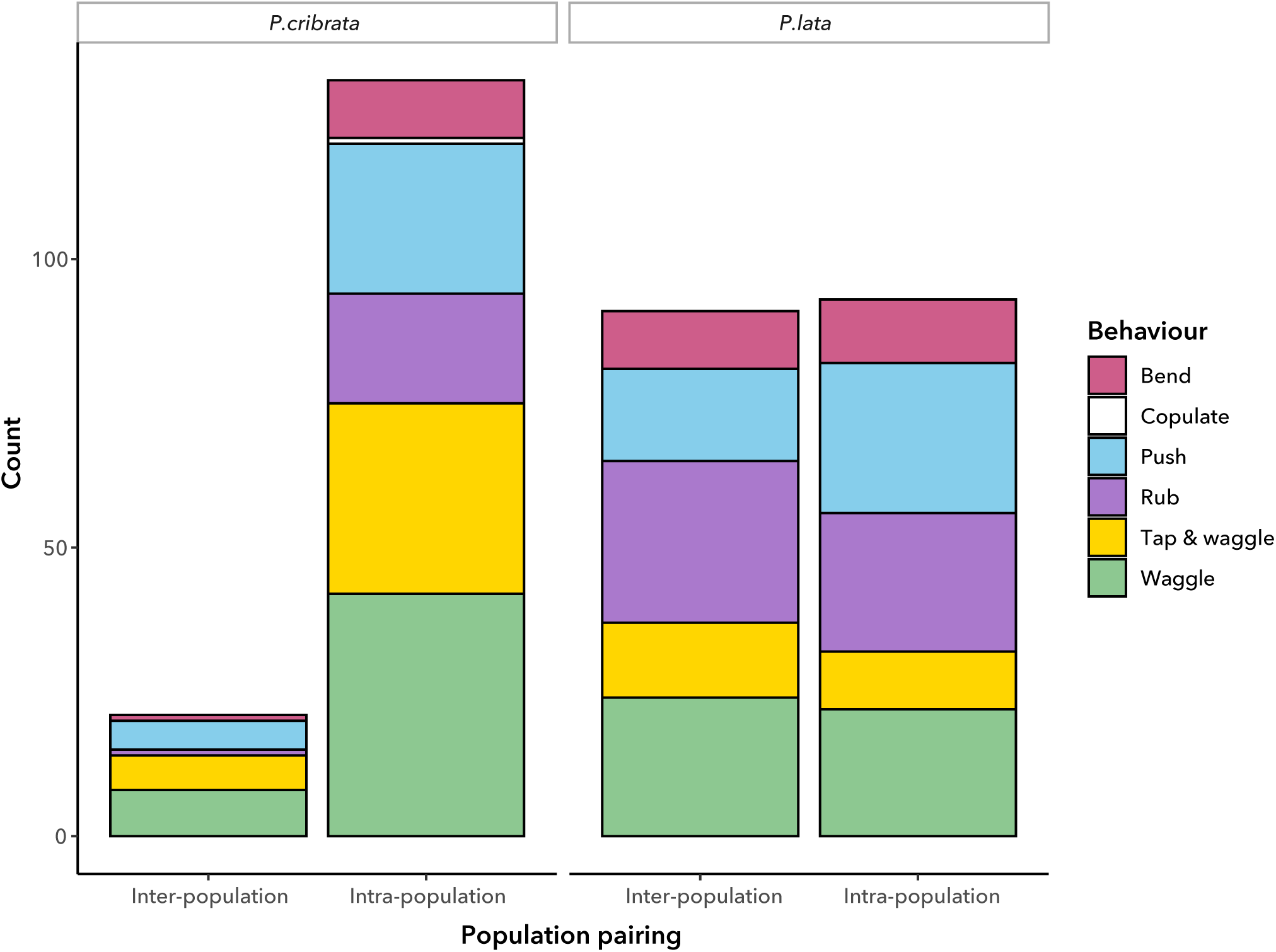
Counts of individual behaviours in courtship interactions, based on observations of male-female pairings.

**Figure 8.**
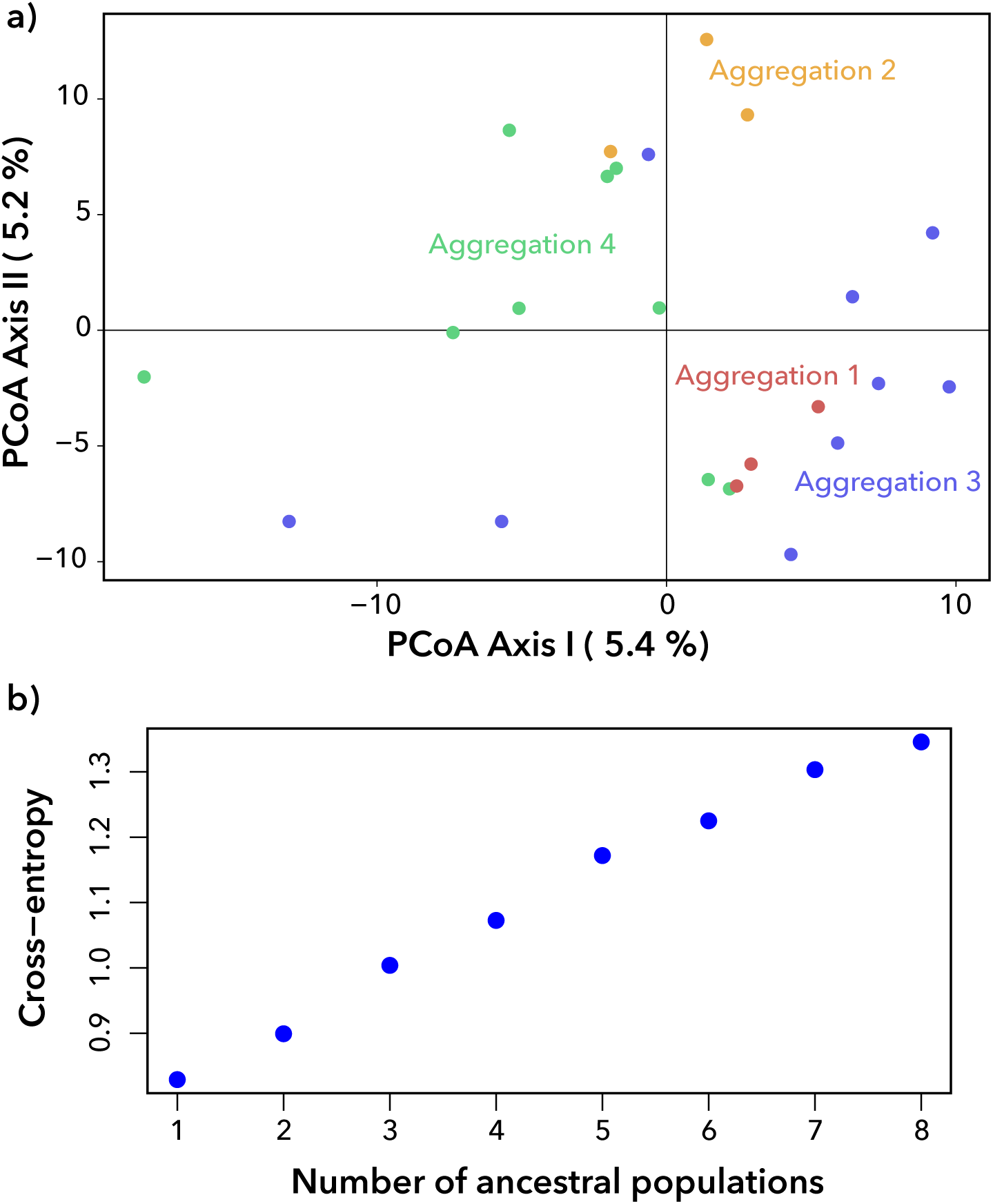
Genetic structure across four nesting aggregations of *Panesthia lata* on Blackburn Island. **a)** PCoA results based on 8,926 nuclear SNPs. **b)** Cross-entropy scores for *K* = 1–8 genetic clusters calculated using sNMF analysis, based on 6,471 nuclear SNPs.

## 4. Discussion

### 4.1. Behavioural shifts following island colonisation

To the best of our knowledge, this study represents the first experimental comparison of behaviour in an island invertebrate with its mainland sister species. We demonstrate that *P. lata* differs from *P. cribrata* along several ecological and behavioural axes, and that these differences are conserved among its different island populations. Most strikingly, male *P. lata* were found to engage in fewer, and less intense, agonistic interactions than *P. cribrata*. While the average duration of interactions was longer in *P. lata*, this appears to be caused by the lower intensity of agonism, which delayed the submission and retreat of the losing individual (S. Coady, pers. obs.). Male agonism in *Panesthia* relates to competition during dispersal and mate-finding (Bell et al., 2007; O’Neill et al., 1987; Rugg & Rose, 1984; Seelinger & Seelinger, 1983), hence our findings suggest a shift in intraspecific competitive dynamics on the LHIG. Due to limitations on the collection of the critically endangered *P. lata*, we acknowledge that the sample size places bounds on interpretation. Nonetheless, the consistency of results across different behavioural metrics, and different agonistic behaviours, underscores their robustness.

Reduced intra-specific aggression is a fundamental pattern in vertebrate island ecology. Such behavioural shifts have been observed across a broad range of organisms that includes mammals (Adler & Levins, 1994; Goltsman et al., 2005), reptiles (Pafilis et al., 2011; Pafilis et al., 2009; Raia et al., 2010) and birds (Jezierski et al., 2023). Two leading hypotheses suggest that this trend is tied to higher population densities on islands, which are linked to lower intraspecific agonism either due to greater resource availability, or because the higher frequency of conspecific interactions elevates energetic and fitness costs of agonism (MacArthur et al., 1972; Stamps & Buechner, 1985; Whittaker et al., 2023; Wright, 1980). It is presently unclear which of these, if any, applies to *P. lata*. We find equivocal evidence of larger aggregation sizes (as a proxy for population density) in *P lata* than *P. cribrata*, although the number of aggregations per unit area appears larger on the LHIG than mainland Australia. Moreover, without comparisons across multiple generations, we cannot presently discern whether the behavioural differences represent plastic responses to differing ecological conditions, or whether they represent fixed, evolved differences. Long-term culture of *Panesthia*, coupled with granular genomic comparisons, will be necessary to untangle the genetic underpinnings.

In contrast to agonistic interactions, we detected no significant differences in the overall frequency or duration of courtship between *P. lata* and *P. cribrata*, and only slight variation in the frequencies of individual behaviours. Courtship behaviour is highly conserved across the Blattodea, which display only four mating strategies (Lizée et al., 2017; Sreng, 1993).

Courtship has also been overlooked in comparisons of island and mainland taxa, although reproductive studies suggest that islands species produce fewer and/or smaller clutches (reviewed by Baeckens & Van Damme, 2020; Novosolov & Meiri, 2013). Due to the long life cycles of *P. lata* and *P. cribrata* (multiple years to reach maturity; Rugg & Rose, 1990), investigation of clutch size was beyond the scope of this project.

One curious point of contrast between the species was the occurrence of male-male courtship in *P. lata*. It is unclear whether this represents a consequence of island existence, as same-sex courtship has been documented across Blattodea (Simon & Barth, 1977). Hypothesised causes including the elevation of sexual excitation, suppression of interspecific hybridisation, and maintenance of courting groups. Same-sex behaviour can also be stimulated by breakdowns in chemical signalling, whereby males are less able to distinguish between sexually receptive and unreceptive females, leading them to court any individuals resembling a conspecific female (Dukas et al., 2006). The high rates of courtship relative to copulation observed presently suggest that male *Panesthia* attempt courtship regardless of female receptivity, which may encourage same-sex courtship (although this was not observed in *P. cribrata*). Comparisons of both same-sex courtship and clutch size in *P. lata*, relative to additional mainland *Panesthia* species, present interesting directions for future study.

### 4.2. Genetic structure of aggregations in P. lata

Surprisingly, SNP analyses uniformly indicated that aggregations on Blackburn Island are composed of unrelated individuals. This sharply contrasts with other *Panesthia* species from Australia and Asia, in which aggregations are thought to primarily consist of parents and offspring (Ito & Osawa, 2019, 2021; Rugg & Rose, 1984). However, this result is consistent with the reduced intraspecific agonism in *P. lata* relative to *P. cribrata*. It also reflects our ecological observations, which suggest that individuals move between stones as frequently as daily.

The evolutionary drivers of this divergent nesting strategy are not clear. The wooden galleries occupied by *P. cribrata* provide both food and shelter, with little benefit from movement between microhabitats. In contrast, *P. lata* forages beyond resident stones or logs (Carlile et al., 2018), which may encourage migration between family groups. We also note that stones are relatively scarce below the *Ficus* tree on Blackburn Island, which could drive unrelated individuals to share shelters. The Australian mainland species *Panesthia matthewsi* is known to occupy similar microhabitats to *P. lata* (shallow burrows below stones). A comparison of aggregation behaviour in *P. lata* and *P. matthewsi* would reveal whether the shift away from obligate log-boring is consistently associated with changes in aggregation structure. Alternatively, the apparent permeability of the aggregations may be tied to inbreeding. Background levels of kinship on Blackburn Island were substantially elevated, and aggregation preferences in *Panesthia* are based in part on chemical cues that distinguish relatives from non-relatives (Billingham et al., 2010). Thus, inbreeding in *P. lata* may interfere with the maintenance of family groups. This latter hypothesis may be examined with behavioural experiments examining association preferences between known close relatives and unrelated individuals.

The island syndrome does not circumscribe a consistent suite of adaptations present in all species, but rather a loose collection of evolutionary tendencies (Jezierski et al., 2023). In line with this expectation, our behavioural and genetic analyses reveal changes in agonism and aggregation behaviour, but not necessarily reproduction, in *P. lata*. Importantly, these changes also appear linked to the unique conditions of insular habitats (Novosolov et al., 2012; Whittaker et al., 2023). Namely, the reductions in agonism and disruption of kin groups may be linked to the higher population densities and/or intensified inbreeding in *P. lata*. It is unclear why invertebrates have been previously overlooked in studies of island behaviour (Gavriilidi et al., 2022), as we demonstrate that *P. lata* is an informative model of island syndrome. Nonetheless, adaptations to islands vary greatly even in established model groups such as passerine birds and rodents (reviewed by Novosolov et al., 2012), thus we emphasise the need for further research across a wide range of invertebrate groups.

### 4.3. Inter-population differences and conservation of P. lata

Finally, in addition to our examination of interspecific behavioural differences, we also undertook morphological and ecological comparisons between populations of *P. lata*. We identified appreciable differences in body length, whereby the North Bay and North Head individuals were smaller and displayed more pronounced sexual size dimorphism (SSD) than that on Blackburn Island. Comparisons to *P. cribrata* suggest that there have been both increases in female size on Blackburn Island and reductions in male body length on LHI. Blackburn Island and LHI have only recently been isolated (*ca.* 10–100 ka; Adams et al., 2025b), however body size is relatively labile within *Panesthia* (based on morphological differences between populations on Blackburn and Roach Islands; H.A. Rose, pers. obs.).

Islands are thought to select for greater SSD, due to relaxed interspecific competition and intensified intraspecific competition (Blanckenhorn, 2005; Meiri et al., 2014). Neither factor clearly explains the divergent morphological trajectories on Blackburn Island and LHI, as our behavioural assays found no difference in agonism across populations, and habitats on both islands appear qualitatively similar (H.A. Rose, pers. obs.). An alternative possibility is that the reduction in male body size may reflect predation from black rats on LHI (e.g. Benítez-López *et al*., 2021), which were historically absent on Blackburn Island. Presuming that male *P. lata* disperse more actively (as in other *Panesthia*; Rugg & Rose, 1984), then males may have experienced stronger predation pressure and selection for smaller size. Museum specimens of *P. lata* collected from Blackburn Island and LHI prior to rodent introduction show no obvious SSD, albeit with a modest sample size (*n* < 20; M.W.D. Adams, pers. obs.).

Despite the morphological differences, we found that neither courtship nor agonistic behaviour varied between inter- and intra-population assays. This is promising for the success of any future conservation translocation between islands: current guidelines recommend evidence of both genetic and ecological compatibility prior to translocation (Hedrick & Fredrickson, 2010). While the present results should be further developed with captive-crossing studies (Stringer & Chappell, 2008), they nonetheless offer preliminary evidence that translocations between North Head and North Bay, or even Blackburn Island, would not disrupt the species’ reproductive ecology.

### 4.4. Conclusions

Here we have presented the first experimental comparison of an island-mainland pair of invertebrate species. Our findings of reduced intraspecific agonism, increased population density and more permeable aggregations in *P. lata* align with ecological and behavioural adaptations seen in island vertebrates. On the other hand, while the reproductive ecology of *P. lata* requires more detailed examination in future, we found no evidence of shifts in courtship behaviour relative to *P. cribrata*. Overall, we provide novel evidence that insect behaviour follows somewhat similar trajectories to vertebrates in island ecosystems.

## Data availability

Raw data and R scripts used in this manuscript are available on Dryad: http://datadryad.org/share/Y0WDIEiqMm7QpWxMgC4uSf9ilwvPwBH1UhaxLK59qwM

## Acknowledgments

This work was supported by the Australia Pacific Science Foundation (grant number: APSF22029) and samples were collected under permits from NSW National Parks and Wildlife Services (SL102663) and the Lord Howe Island Board (05/ 22). We thank Toby G.L. Kovacs, Campbell Wilson, Younis Menkara, Simon Y.W. Ho and Chris A.M. Reid for assistance during fieldwork, the Lord Howe Island Board for logistical support, and Brendan Coady for constructing the cockroach arenas. The authors declare no conflicts of interest.

