## Supplementary Material for "Island syndrome in the critically endangered Lord Howe Island cockroach *Panesthia lata*"

### Supplementary Results

*Ecological observations of the North Bay population*

During the survey of the North Bay population on May 21st, 2023, 13 adult females, 11 adult males and 92 nymphs were found across two 45-degree segments examined under the banyan tree (one N to NW, and one S to SE), with the assistance of Toby Kovacs (University of Sydney). Individuals were found as far as 12 m from the central trunk of the tree, however only two nymphs were found beyond the canopy cover of the banyan tree. Our most conservative population estimate of the North Bay population was >400 individuals total, including *ca.*100 adults. In each segment there were some rocks that were too large to move, as well as areas with large abundant large roots that could not be easily surveyed for cockroaches. Additionally, once a rock was lifted, individual *P. lata* would bury themselves and become very difficult to spot. It is likely that some individuals were buried and remained uncounted.

*Ecological observations of the North Head population*

Chris Reid (Australian Museum) and NL examined *P. lata* below the canopy of a single *F. macrophylla* tree near the top of the ridge of North Head at ~10 am on Thursday 20th February, 2025. Cockroaches were detected by overturning stones and logs along a single northwest transect extending from the tree trunk to the margin of the tree canopy (approximately 18 m in length). We recorded 32 adults and 80 nymphs ($n$ = 112 in total) under a total of 17 rocks, leading to a coarse estimated population size of 896 individuals. Due to the cryptic nature of the species, and the presence of several large rocks that could not be overturned in our surveys, we consider this number to be a minimum bound on the population size. Previous examinations of the ground under three banyan other trees along the North Head ridgeline on May 24th, 2023 by Toby Kovacs, SC, and NL revealed they were covered in a thick invasive vine, making it difficult to turn over rocks underneath the trees. No overturned rocks showed signs of *P. lata*.

Following the return of the material to Sydney, a number of first-instar nymphs were found in culture, indicating that females can give birth under laboratory conditions. Whether the births were due to copulations that occurred in the wild, or in the laboratory, is unknown.

*SNP generation*

**Supplementary Table S1.** Samples of *Panesthia lata* used in genetic analyses. Abbreviations: Australian Museum (AM), Australian National Insect Collection (ANIC), H.A. Rose private collection (HARPC), single-nucleotide polymorphism (SNP). Missing data quantified prior to filtering.

| Sample ID | Aggregation | Locality | Collection institution | Year collected | DNA extraction method | SNP missing data (%) |
|---|---|---|---|---|---|---|
| PL3 | – | Blackburn Island | HARPC | 2022 | ANIC | 21.2 |
| PL4 | – | Blackburn Island | HARPC | 2022 | ANIC | 21.0 |
| PL6 | – | Blackburn Island | HARPC | 2022 | ANIC | 19.2 |
| PL7 | – | Blackburn Island | HARPC | 2022 | ANIC | 21.6 |
| PL17 | – | Blackburn Island | HARPC | 2022 | In-house | 21.8 |
| PL33 | – | Blackburn Island | HARPC | 2022 | In-house | 18.9 |
| PL34 | – | Blackburn Island | HARPC | 2022 | In-house | 18.1 |
| PL35 | – | Blackburn Island | HARPC | 2022 | In-house | 18.4 |
| PL36 | – | Blackburn Island | HARPC | 2022 | In-house | 18.4 |
| PL38 | – | Blackburn Island | HARPC | 2022 | In-house | 19.3 |
| PL41 | – | Blackburn Island | HARPC | 2022 | In-house | 19.2 |
| PL42 | 1 | Blackburn Island | HARPC | 2022 | In-house | 18.3 |
| PL43 | 1 | Blackburn Island | HARPC | 2022 | In-house | 19.0 |
| PL44 | 1 | Blackburn Island | HARPC | 2022 | In-house | 18.7 |
| PL45 | 2 | Blackburn Island | HARPC | 2022 | In-house | 18.6 |
| PL46 | 2 | Blackburn Island | HARPC | 2022 | In-house | 18.5 |
| PL47 | 2 | Blackburn Island | HARPC | 2022 | In-house | 21.5 |
| PL48 | 3 | Blackburn Island | HARPC | 2022 | In-house | 21.0 |
| PL49 | 3 | Blackburn Island | HARPC | 2022 | In-house | 21.1 |
| PL50 | 3 | Blackburn Island | HARPC | 2022 | In-house | 20.9 |
| PL51 | 3 | Blackburn Island | HARPC | 2022 | In-house | 20.6 |
| PL52 | 3 | Blackburn Island | HARPC | 2022 | In-house | 20.8 |
| PL53 | 3 | Blackburn Island | HARPC | 2022 | In-house | 21.8 |
| PL54 | 3 | Blackburn Island | HARPC | 2022 | In-house | 21.9 |
| PL55 | 3 | Blackburn Island | HARPC | 2022 | In-house | 19.4 |
| PL56 | 3 | Blackburn Island | HARPC | 2022 | In-house | 19.4 |

| | | | | | | |
|---|---|---|---|---|---|---|
| PL57 | 4 | Blackburn Island | HARPC | 2022 | In-house | 18.8 |
| PL58 | 4 | Blackburn Island | HARPC | 2022 | In-house | 19.8 |
| PL59 | 4 | Blackburn Island | HARPC | 2022 | In-house | 19.3 |
| PL60 | 4 | Blackburn Island | HARPC | 2022 | In-house | 19.6 |
| PL61 | 4 | Blackburn Island | HARPC | 2022 | In-house | 18.6 |
| PL62 | 4 | Blackburn Island | HARPC | 2022 | In-house | 19.5 |
| PL63 | 4 | Blackburn Island | HARPC | 2022 | In-house | 21.6 |
| PL64 | 4 | Blackburn Island | HARPC | 2022 | In-house | 21.2 |
| PL65 | 4 | Blackburn Island | HARPC | 2022 | In-house | 20.4 |
| PL1 | – | North Bay | HARPC | 2022 | ANIC | 22.8 |
| PL2 | – | North Bay | HARPC | 2022 | In-house | 21.6 |
| PL5 | – | North Bay | HARPC | 2022 | In-house | 27.4 |
| PL18 | – | North Bay | HARPC | 2022 | In-house | 25.6 |
| PL19 | – | North Bay | HARPC | 2022 | In-house | 23.7 |
| PL26 | – | North Bay | HARPC | 2022 | In-house | 21.4 |
| PL27 | – | North Bay | HARPC | 2022 | ANIC | 22.2 |
| PL28 | – | North Bay | HARPC | 2022 | ANIC | 21.8 |
| PL29 | – | North Bay | HARPC | 2022 | In-house | 21.4 |
| PL30 | – | North Bay | HARPC | 2022 | In-house | 21.2 |
| K.383796 | – | Roach Island | AM | 2000 | ANIC | 27.2 |
| K.383797 | – | Roach Island | AM | 2000 | In-house | 88.4 |
| K.383798 | – | Roach Island | AM | 2000 | ANIC | 48.9 |
| K.383799 | – | Roach Island | AM | 2000 | ANIC | 41.2 |
| K.383800 | – | Roach Island | AM | 2000 | ANIC | 82.9 |
| K.383801 | – | Roach Island | AM | 2000 | In-house | 23.7 |
| PL8 | – | Roach Island | HARPC | 2003 | In-house | 34.0 |
| PL9 | – | Roach Island | HARPC | 2003 | In-house | 33.4 |
| PL10 | – | Roach Island | HARPC | 2003 | In-house | 29.4 |
| PL11 | – | Roach Island | HARPC | 2003 | In-house | 25.7 |
| PL12 | – | Roach Island | HARPC | 2003 | In-house | 22.8 |
| PL13 | – | Roach Island | HARPC | 2003 | In-house | 22.7 |
| PL14 | – | Roach Island | HARPC | 2003 | In-house | 26.3 |

| | | | | | | |
|---|---|---|---|---|---|---|
| PL15 | – | Roach Island | HARPC | 2003 | In-house | 26.3 |
| PL16 | – | Roach Island | HARPC | 2003 | In-house | 24.1 |
| PL20 | – | Roach Island | HARPC | 2003 | In-house | 99.7 |
| PL21 | – | Roach Island | HARPC | 2022 | In-house | 25.2 |
| PL22 | – | Roach Island | HARPC | 2022 | In-house | 21.8 |
| PL23 | – | Roach Island | HARPC | 2022 | In-house | 21.7 |
| PL24 | – | Roach Island | HARPC | 2022 | In-house | 22.4 |
| PL25 | – | Roach Island | HARPC | 2022 | In-house | 22.8 |
| PL66 | – | Roach Island | HARPC | 2022 | In-house | 35.7 |

### Results

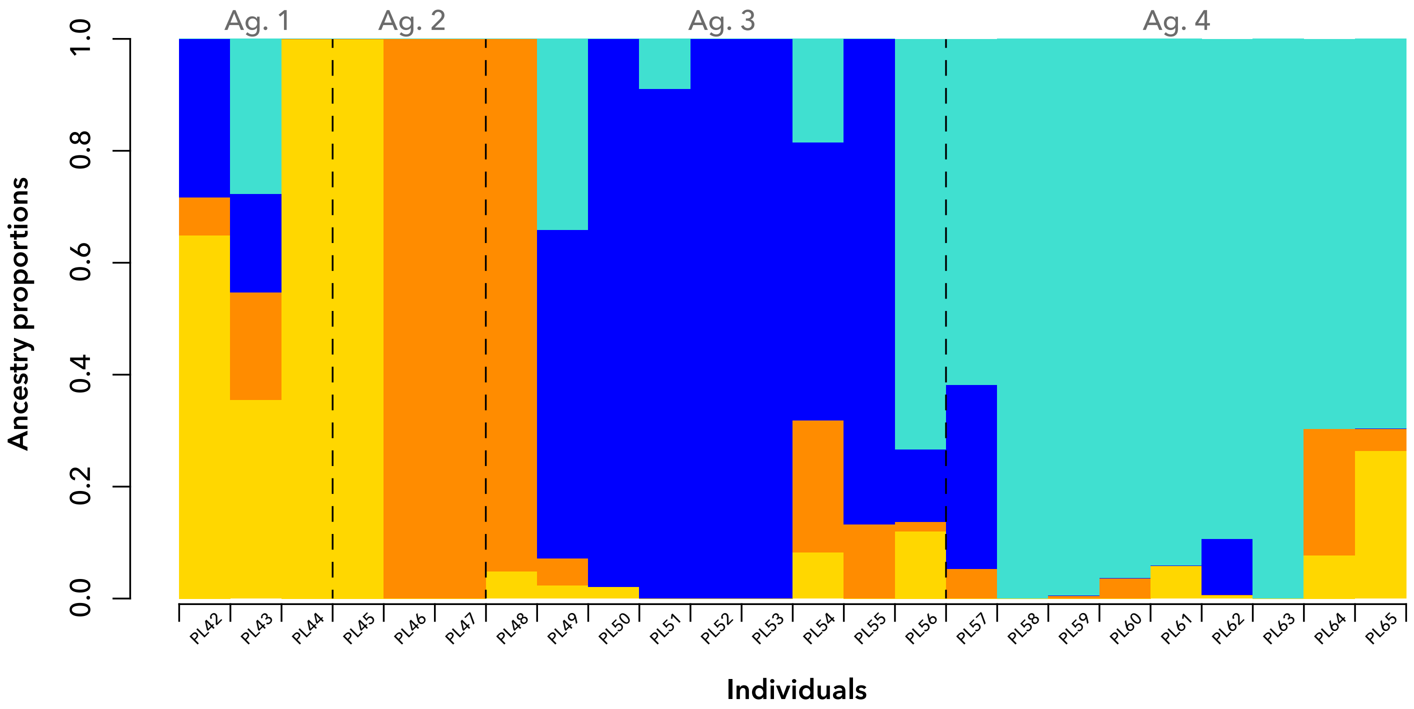


**Supplementary Figure S1.** Admixture plot for four aggregations of *Panesthia lata* collected from separate rocks on Blackburn Island with $K = 4$. Ancestry estimated using a sparse non-negative matrix factorization algorithm in *LEA*. Estimated genetic clusters do not align with any of the four aggregations (ag. 1–4, denoted by dashed lines).
